# ATAT: Automated Tissue Alignment and Traversal in Spatial Transcriptomics with Self-Supervised Learning

**DOI:** 10.1101/2023.12.08.570839

**Authors:** Steven Song, Emaan Mohsin, Renyu Zhang, Andrey Kuznetsov, Le Shen, Robert L. Grossman, Christopher R. Weber, Aly A. Khan

## Abstract

Spatial transcriptomics (ST) has enhanced RNA analysis in tissue biopsies, but interpreting these data is challenging without expert input. We present Automated Tissue Alignment and Traversal (ATAT), a novel computational framework designed to enhance ST analysis in the context of multiple and complex tissue architectures and morphologies, such as those found in biopsies of the gastrointestinal tract. ATAT utilizes self-supervised contrastive learning on hematoxylin and eosin (H&E) stained images to automate the alignment and traversal of ST data. This approach addresses a critical gap in current ST analysis methodologies, which rely heavily on manual annotation and pathologist expertise to delineate regions of interest for accurate gene expression modeling. Our framework not only streamlines the alignment of multiple ST samples, but also demonstrates robustness in modeling gene expression transitions across specific regions. Additionally, we highlight the ability of ATAT to traverse complex tissue topologies in real-world cases from various individuals and conditions. Our method successfully elucidates differences in immune infiltration patterns across the intestinal wall, enabling the modeling of transcriptional changes across histological layers. We show that ATAT achieves comparable performance to the state-of-the-art method, while alleviating the burden of manual annotation and enabling alignment of tissue samples with complex morphologies.

**Availability:** ATAT is available at: https://github.com/StevenSong/tissue-alignment

## 1 Introduction

Spatial transcriptomics (ST) has become an increasingly utilized and powerful tool for understanding the cellular landscape within tissues [1]. By merging spatial gene expression profiling with histological imaging, it offers a comprehensive view of tissue architecture and function. While histological images are crucial for comparing pathology across various cases and conditions, we have yet to fully explore their capacity to aid in comparing gene expression across diverse spatial transcriptomics datasets [2, 3]. Leveraging this information could establish a common histologically-defined axis for spatial gene expression data, addressing the crucial challenge of comparing gene expression across tissue samples in a way that respects anatomical and cellular contexts.

Spatially resolved transcriptomics often generates two types of overlaid data: gene expression data and histological images. This overlay provides a composite view of tissue architecture and function, crucial for understanding phenomena such as drug responses or disease pathology. A case in point is the variability in immune cell infiltration depth within the intestinal mucosa, which can have implications for the severity of inflammatory conditions and can vary depending on disease etiology [4]. In order to identify tissue layers, e.g. to determine the depth of inflammation, ST data can be manually annotated and aligned by pathologists. Yet this is a labor intensive and expensive task that can be prohibitive for the wider adoption of ST. Existing computational methods have largely relied on gene expression information to guide the delineation of specific histological layers [5, 6, 7, 8, 9]. Although one could argue that tissue morphology is intrinsically linked to its transcriptional profile [10], the use of gene expression both to first define layers and to then establish transcriptional gradients introduces a circular bias.

As such, a predominant limitation in the current ST landscape is an overemphasis on expression information without adequately considering the invaluable insights provided by histological images about tissue morphology. Tissues are inherently organized into histological layers, each possessing its unique cellular architecture. By understanding this layered arrangement, it becomes possible to project expression data from particular layers along a one-dimensional histologically-defined axis. We term this projected expression as a *gene expression trajectory*. This organized representation can facilitate straightforward comparisons of gene expression across similar tissue layers/axes between different samples. For this reason, gene expression trajectories are an important end point for modern ST analysis methods [5]. However, the absence of pathologist-marked regions in datasets often complicates modeling gene expression over exact histological layers [11, 12], highlighting the need for automated methods that capture essential spatial information from histological imaging [2]. Current methods that incorporate histology images often resort to naive measures, such as averaging colors across tiles, which can potentially miss out on key cellular morphology nuances [13]. Other methods have integrated expression data with image features from pre-trained representations that are derived from non-histological images [14]. Although the field of digital pathology is distinct from ST, it has experienced a resurgence due to the integration of computer vision techniques. Recent advancements leverage self-supervised learning to discern contextual domains within images, aiding in the identification of specific histological layers in H&E images, such as the mucosa in gut biopsies [15].

A long-standing challenge in the current ST landscape is inter-slide comparison, especially across different tissue samples or conditions [2, 3]. Current methods primarily focus on serial sections for slide comparison [16]. At the same time, expression-based methods struggle to reliably overlay distinct tissue specimens [17, 18]. A practical approach is to use consistent anatomical and histological structures present in tissues as anchoring reference points. Definitions of layers based on gene expression can be imprecise due to overlaps in transitional zones, discrepancies across samples both within a disease and across different diseases, or changes induced by inflammation and disease. These structures, clearly depicted in histology images, can help standardize slide alignment. These structures can be directly inferred from the histological imaging data, thus offering an unbiased method to define structure and align across multiple tissue samples, breaking the circular reasoning present in expression-only based methods.

Using histology images for aligning histological layers brings dual advantages. By focusing on universally recognizable histological landmarks, this strategy avoids relying on condition specific gene expression profiles. It also offers a method independent of the often sparse and variable ST data [19]. Prioritizing image-centric alignment avoids the pitfalls of depending solely on spatially resolved transcriptomics data, data which is vulnerable to numerous factors including technical anomalies, inherent biological conditions, and sparsity.

Leveraging histological structures for unbiased alignment allows for genuine evaluation of gene expression differences across conditions. For example, it becomes feasible to dissect how a specific disease influences gene expression throughout tissue layers when these layers are consistently defined. Adopting this approach not only strengthens analysis of ST but also maximizes the transformative potential of ST, revealing deep insights about cellular dynamics and tissue functions in both health and disease.

**Our contribution** In this paper, we introduce ATAT: Automated Tissue Alignment and Traversal, an algorithm which, to our knowledge, is the first algorithm that utilizes self-supervised contrastive learning over the H&E image to align and traverse ST data. We propose a surprisingly simple and efficient integrated algorithmic framework that is tailored to the dataset and utilizes the H&E images of a tissue biopsy to automatically chart a path across a sample with any morphology. We generated ST data from different individuals and conditions, which brought to light the complex topology of tissue sections present in real world cases. We applied our method to successfully elucidate differences in immune infiltration patterns across the intestinal wall. This enables us to model transcriptional changes across histological layers, thereby reducing the burden of manual segmentation of tissue samples and complementing expression-only based methods.

## 2 Methods

### 2.1 Data collection and preprocessing

We developed ATAT on tissue samples collected from 12 colon and 4 stomach resections under protocol IRB21-0929 approved by the University of Chicago Institutional Review Board. Of the 12 colon sections, 4 are from patients with ulcerative colitis (UC), 4 are from patients with Crohn’s disease (CD), 3 are from patients with *Clostridoides difficile* (C. diff) infection, and 1 is from a patient with normal colon. Of the 4 stomach sections, 3 are from patients with *Helicobacter pylori* (H. pylori) infection and 1 is from a patient with normal stomach. Each tissue sample was processed using the 10x Genomics Visium Spatial Gene Expression platform. Sequencing data was preprocessed using the 10x Genomics Space Ranger software v1.3.1. We also apply ATAT to 12 tissue samples from the dorsolateral prefrontal cortex (DLPFC) [20]. For each H&E stained slide, the ST data is resolved as a hexagonal grid of spots. Within the square 6.5mm tissue capture area, there are 78 rows and 64 columns of spots, with each spot covering an area 55μm in diameter, though the exact number of spots depends on the size of the tissue sample.

### 2.2 Gene expression normalization

We normalize our ST data per slide. Given a gene expression matrix, *X*, we normalize the nts n each spot (i.e. tile), similar to normalization of single-cell RNA seq counts [21]: 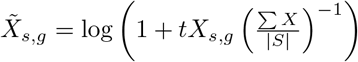, where 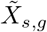 is the normalized count of gene *g* at spot *s, t* is the target average count per spot (we set *t* = 10^4^), *S* is the set of all spots, and 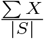 represents the average count per spot.

### 2.3 Automated tissue alignment and traversal

To plot ST data across tissue layers, we first learn a spatially aware representation of each image tile centered on the ST spots. Next, we calculate a similarity score for every pair of adjacent tiles. Using these scores as the weights for the edges, we identify the shortest path between a user-specified starting point and endpoint on the slide. Finally, we assign each tile on the slide to the tiles along the path and average the gene expression along the path of all tiles with the same assignment. Figure 2 depicts a graphical overview of ATAT and the pseudocode for ATAT is described in Algorithm 1.

**Figure 1:**
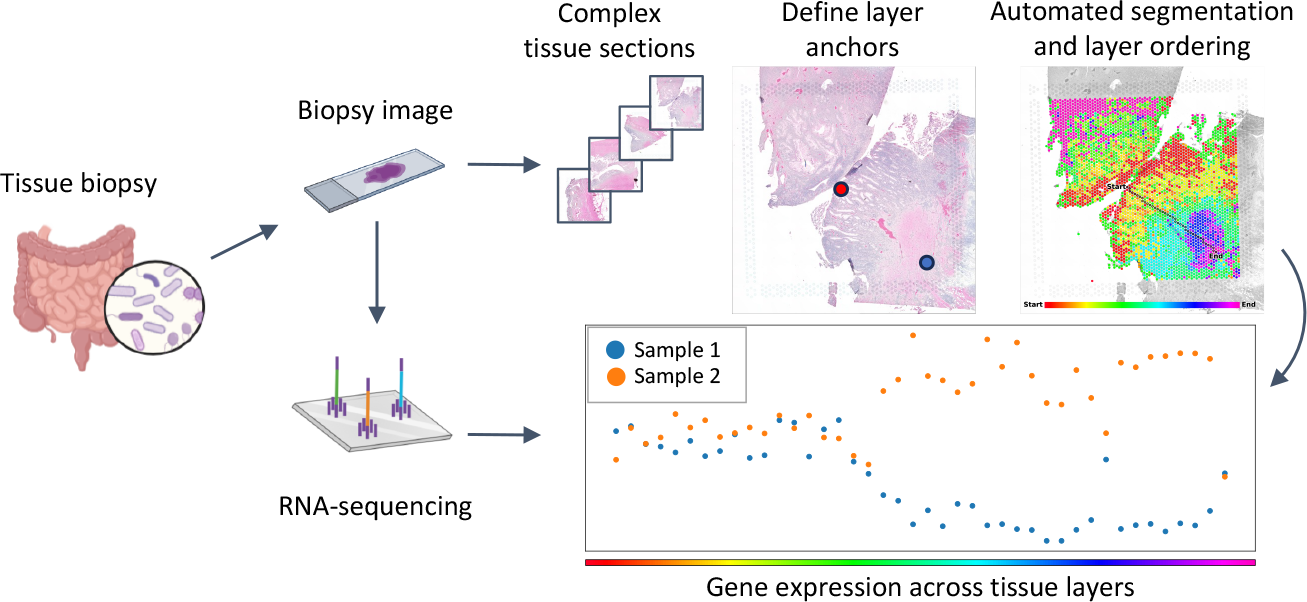
Overview of the ATAT workflow. Tissue samples are imaged and sequenced using ST methods. Many of these sections have complex tissue morphologies. ATAT utilizes the H&E image to traverse tissue layers between biological anchor points. These anchored paths allow for comparisons of gene expression trajectories between multiple tissue samples with similarly anchored paths.

**Figure 2:**
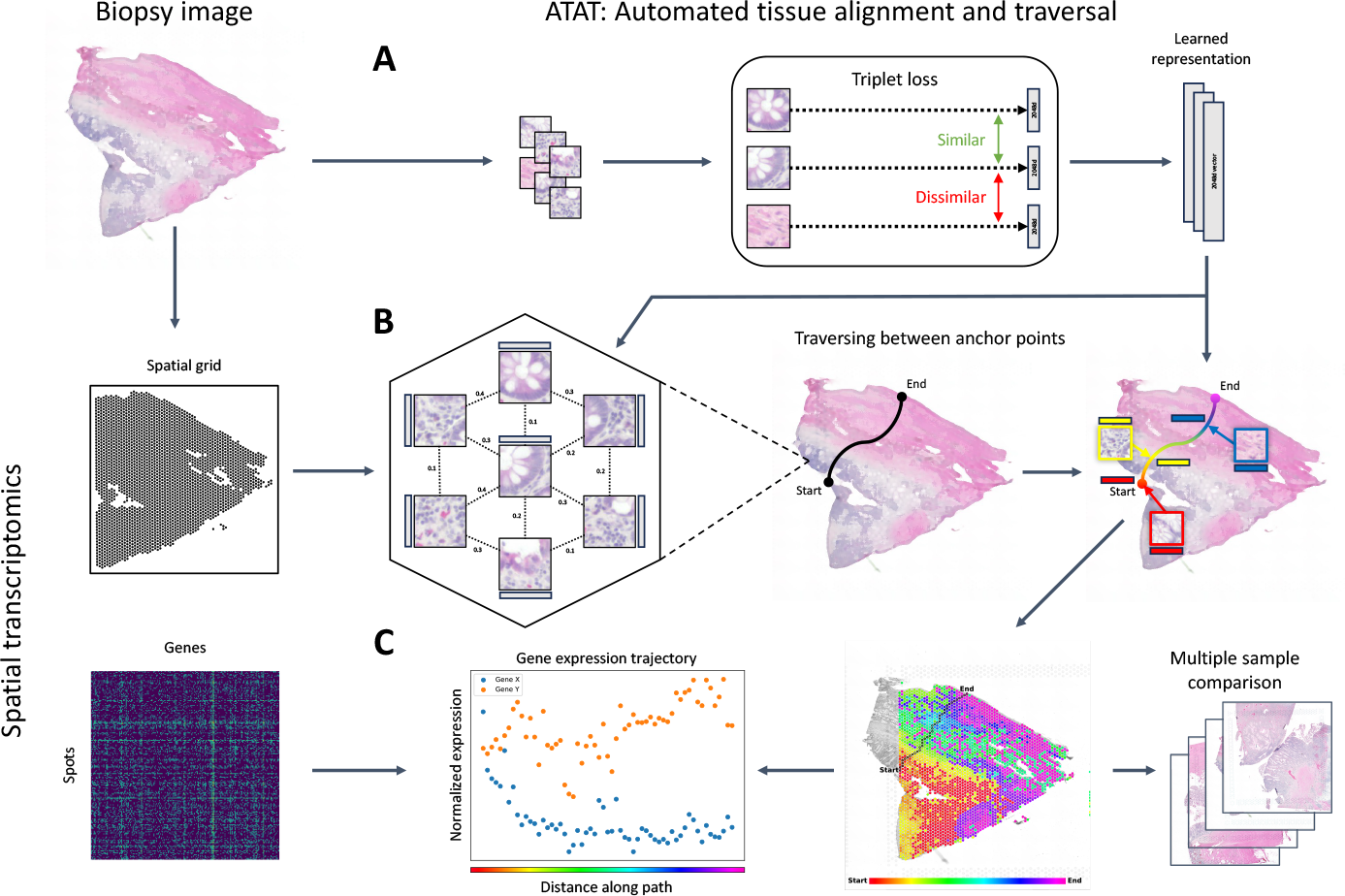
An overview of the Automated Tissue Alignment and Traversal (ATAT) algorithm. (A) Learning spatially aware tile representations. A tissue sample H&E slide is cut into tiles and used to train a convolutional neural network using triplet loss to learn visually similar tiles and gradual transitions between adjacent tiles. (B) Path traversal through the lattice structured graph. Between adjacent tiles on the spatial grid, a similarity score is calculated using the learned tile representations. A path is traversed between user selected start and end anchor points using the similarity score as edge weights for a shortest path algorithm between adjacent tiles. The similarity between other tiles on the slide and all tiles on the path are used to assign sets of tiles across the slide to tiles on the path. (C) Deriving gene expression trajectories along traversed paths. The gene expression at each tile along the path is averaged across the set of all tiles assigned to the path tile. Comparisons of gene expression patterns across multiple samples is enabled by path traversal through similarly anchored paths.

#### 2.3.1 Learning spatially aware tile representations

We first extract tiles from H&E slides. Each tile is a square with side length equal to the spot diameter in pixels and centered on the spots in the hexagonal grid. We normalize each H&E tile by first subtracting, per channel, the average of all tiles in the dataset. We then divide by, per channel, the standard deviation of all tiles in the dataset.

We train a convolutional neural network (CNN) using triplet loss over the dataset to learn a spatially aware tile representation (Figure 2A). For the CNN, we utilize a resnet50 architecture with a 3 layer multilayer perceptron on top with hidden dimension 2048, as in Chen and He [22]. For our experiments, we train the model for 1000 epochs with batch size 768, learning rate 0.05, and SGD optimizer with momentum 0.9 and weight decay 1e-4.

Triplet loss is a margin-based loss function used to learn useful embeddings by distance metric learning. We employ triplet loss to cluster tissue tiles into distinct histological regions, ensuring that tiles of similar histological types are pulled closer while those of different types are pushed apart in the embedding space [15]. Formally, given a triplet of tiles ⟨*a, p, n*⟩ where *a* is an anchor tile, *p* is a positive tile randomly sampled from the tiles immediately adjacent to *a*, and *n* is a negative tile randomly sampled from all other tiles in the dataset, the triplet loss is defined as: *L*_triplet_ = max(0, *m*(*f* (*a*), *f* (*p*)) *− m*(*f* (*a*), *f* (*n*)) + *α*), where *m* is a distance metric (we use mean squared error for training), *f* denotes the embedding function learned by the CNN, and *α* is the margin that is enforced between positive and negative pairs (0.001).

#### 2.3.2 Path traversal through the lattice structured graph

After the model is trained with triplet loss, we then leverage the hexagonal lattice structure of the spatial profiling array to define a graph (*V, E*), where *V* represents the vertices (tissue tiles) and *E* represents the edges (adjacent tile relationships). We define an adjacency matrix *W* weighted by the distance between tile representations inferred from triplet loss: *W*_*i,j*_ = *m*(*f* (*s*_*i*_), *f* (*s*_*j*_)), where *m* is a distance metric (we use Euclidean distance for the adjacency matrix), and *s*_*i*_, *s*_*j*_ are tissue tiles.

After defining a start vertex *a* and end vertex *b* from user input, we find the shortest path from *a* to *b* through this weighted graph that traverses the histological landscape (Figure 2B); Dijkstra’s algorithm or a similar shortest-path algorithm can be employed to navigate through the lattice structure: 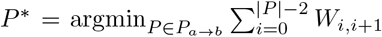 ^|^, where

##### Algorithm 1

ATAT Algorithm

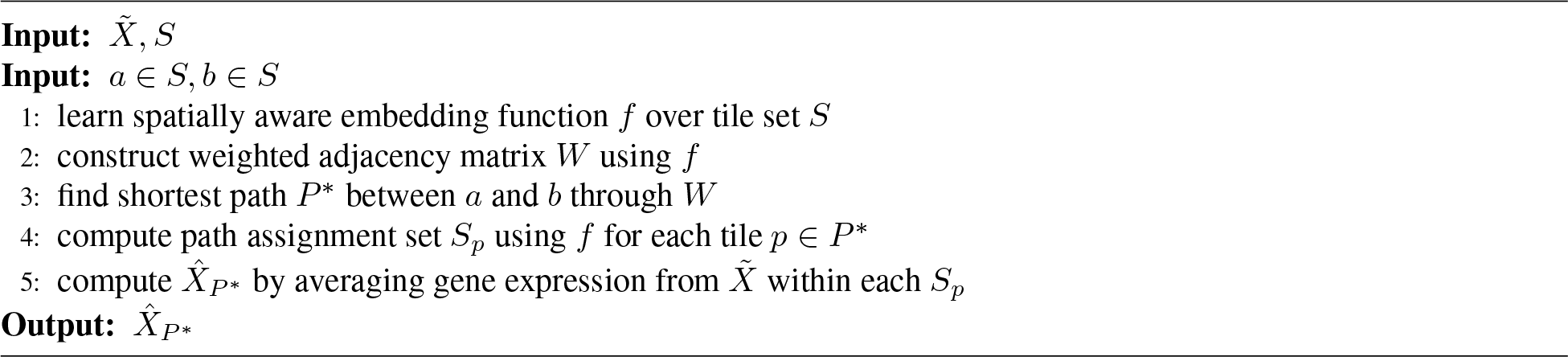

*P*_*a→b*_ denotes the set of all possible paths through the grid from *a* to *b*, and *P* ^*\**^ is the optimal path traversing through the tissue landscape. This path respects spot adjacency and thus does not skip over spots, while also finding the most direct path through tissue layers, following the biological organization of tissue by depth within each layer.

#### 2.3.3 Aligning tiles to the traversed path

Upon determining the optimal path *P* ^*\**^ through the lattice-structured spatial profiling array, we aim to align the rest of the tiles in the slide to the tiles along the path. This is achieved by assigning membership of any tile in the slide to a tile on the path, thereby constructing an assignment set of tiles for each tile along the path. The length of the path signifies the number of assignment sets, | *P* ^*\**^| = *k*. Formally: *S*_*p*_ = *s* |*C*(*s*) = *p*, 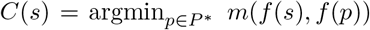, where *S*_*p*_ is the set of tiles *s* assigned to tile 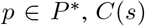 is the assignment for tile *s* and *m* is a distance metric (we also use Euclidean distance for tile alignment). This assignment effectively projects the tissue sample onto the 1-dimensional path utilizing the learned spatial and histological similarity between tiles. With the assignments sets, gene expression can be averaged within each set along the path *P* ^*\**^, resulting in 1-dimensional expression trajectories which can be further used for analysis (Figure 2C). The average gene expression along the path can be expressed as: 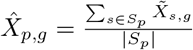, where 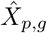is the average gene expression at tile *p ∈ P* ^*\**^ for gene *g, S*_*p*_ is the assignment set for tile *p ∈ P* ^*\**^, and 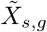 is the normalized gene expression of tile *s* for gene *g*. For convenience, we denote the path aligned gene expression matrix as 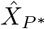, representing the aligned gene expression trajectories for all genes.

### 2.4 Comparing gene expression trajectories from multiple samples

For a given sample, a path traversed by ATAT will likely have a different number of spots *k* which make up the path, as compared to paths traversed through other samples. This results in differing numbers of path tile assignment sets for each path. However, by anchoring the start and end points for each path in the same biological tissue layers across all samples, each of the paths traverse their respective tissue sample in the same order and to a similar depth. Under this assumption, the resulting expression trajectories from alignment can be compared by stacking either end of each sample’s trajectories together. Then the number of assignment sets *k* for a given sample’s path and alignment can be thought of as a reflection of the sampling rate of the trajectory, scaled by the relative amount of time the trajectory spends in each tissue layer.

To achieve this comparison across samples, we assume a relative distance of 0 for the start and 1 for the end of each path, effectively normalizing the length of the path. For each path, we then linearly interpolate the expression trajectory along the path at every 0.01 interval between [0, 1] to generate a new path-length-normalized aligned gene expression matrix, 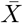. Each 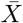 has shape (101, |*G*|), where |*G*| is the set of all genes. We then average or compare length-normalized expression trajectories as outlined below.

### 2.5 Method evaluation

We evaluate our method for internal and external consistency by comparing the resulting gene expression trajectories from multiple alignments of a single tissue sample. For internal consistency, we first establish a set of “reasonable” start/end pairs from which to randomly sample from. For each randomly sampled pair, we traverse and align the tissue sample using ATAT. We additionally align the tissue using an external method which relies on manual segmentation. From each tissue alignment, we generate and length-normalize gene expression trajectories to allow comparisons between trajectories. For these evaluations, we selected UC sample B from Figure 3C due to its relatively simple tissue organization and orientation. Finally, across all CD and UC samples, we evaluate our method’s ability to align tissue by comparing segmentations derived from ATAT to the segmentation from other methods.

**Figure 3:**
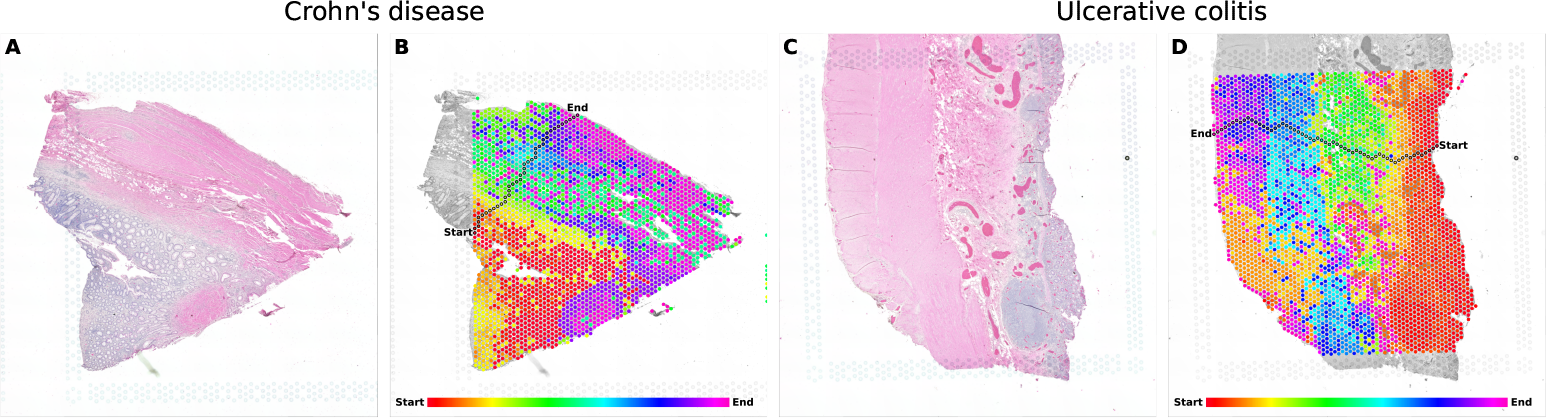
Traversal and alignment of colon tissue samples, anchored by the anatomical layers of the colon. (A) The H&E image of Crohn’s disease sample B. (B) The ATAT alignment of the slide from A overlayed by the hexagonal grid of spots. Between a user selected start and end point, the path traversed by ATAT is outlined in black with each spot along the path colored by a rainbow gradient. The other spots on the slide are shaded by the same color as the spot along the path which is most visually similar. The underlying H&E slide is in grayscale for visual contrast only. (C, D) The same as A and B for ulcerative colitis sample B. The color gradient for B and D are distinctly mapped to their respective path lengths.

#### 2.5.1 Random path sampling

To evaluate if our method is invariant to the selection of start and end spots, we aim to create a set of randomly sampled start and end spots as the basis for our evaluation. We utilize UC sample B for this evaluation. Since we assume a research user would select “reasonable” start and end spots, we create a sampling strategy to mimic the selections a user would make on this sample. At a high level, these “reasonable” start and end pairs help to anchor the traversed path to the actual underlying tissue layers. To define a reasonable start/end pair, the pair should a) be on the edge of the outer tissue layers and b) be connected by a line which is relatively perpendicular to the layer boundaries of the tissue. From the first criterion, we can create a set of potential start spots, *A ⊂ S*, and potential end spots, *B ⊂ S*. For each start spot *a*_*i*_*∈ A*, we find a corresponding set of end spots *B*_*i*_ *⊆ B* which satisfies the second criterion. Then the set of all reasonable start/end pairs is: *Z* = {⟨*a*_*i*_, *b*_*j*_⟩ |*a*_*i*_ *∈A, B*_*i*_ *⊆ B, b*_*j*_ *∈ B*_*i*_, *i <*| *A*|, *j <* |*B*_*i*_|} .

This procedure requires knowing the orientation and architecture of the specific tissue sample. In our evaluation sample, the tissue layers are stacked horizontally with the edges of the outer layers contained within the capture area; the set of potential start or end spots correspond to the right or left edge of the sample, respectively. Additionally, the edges of the outer layers are oriented vertically and are aligned at the top of the slide. We create an ordered set of start spots, *A ⊂ S*, and an ordered set of end spots *B ⊂ S*, sorted in ascending order by the y-coordinate of each spot. Given this ordering, *a*_*i*_ *∈A* corresponds to *b*_*i*_ *∈B*. To construct the set *B*_*i*_ *⊆ B*: *B*_*i*_ = {*b*_*j*_| *∈b*_*j*_ *B, j ≥*max(*i− r*, 0), *j <* min(*i* + *r* + 1, |*B*|), where *r* is the radius on either side of *b*_*i*_ from which to include spots (we set *r* = 15). This results in |*B*_*i*_|*≤* 2*r* + 1. We then construct *Z* as above, resulting in 1, 796 reasonable start/end pairs in our evaluation sample. Finally, we randomly sample without replacement 100 pairs as 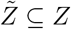. We illustrate this sampling of reasonable paths in Figure S3. We run ATAT to generate aligned gene expression matrix 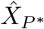 on each randomly sampled start/end pair. Denote the set of these aligned gene expression matrices as 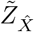 and let 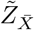be the set of length-normalized matrices from the randomly sampled paths.

#### 2.5.2 Belayer alignment

We compare ATAT against Belayer, a method which currently achieves state-of-the-art performance in aligning tissue samples [5]. We specifically compare against the Belayer mode which takes as input manually annotated layer boundaries and returns a scalar depth value for each spot on the slide. The gene expression is then averaged across all spots of a similar depth. We refer the reader to the original paper for more details on the Belayer method [5].

To run Belayer on our tissue samples, we first manually annotate layer boundaries. If a tissue sample comes with an accompanying layer segmentation, i.e. a layer assignment for each spot on the slide, we still must manually annotate the boundaries of each segment as the relevant boundaries also depend on the orientation and architecture of the sample. Along with the hexagonal layout of the spots, these boundaries are difficult to automatically extract from a true segmentation using simple heuristics. We use the manually annotated layer boundaries as input to Belayer to generate a scalar depth value *d*_*s*_ for each spot *s* on the slide. Then the assignment set for tile *s* is: *C*_Belayer_(*s*) = round(*d*_*s*_).

As before, we average gene expression across all spots within an assignment set (in this case, all spots at a similar depth), resulting in an aligned gene expression matrix 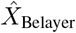. See Section 2.3.3 for more details on gene expression averaging from tile assignments. To length-normalize the expression matrix of Belayer, we use the number of unique rounded depths values as *k* and assume the rounded depths are linearly spaced. We find this approximation is sufficient with almost no missing values between the minimum and maximum depth for our samples, though scaling the interpolation by the actual Belayer depth is also possible. Let 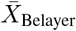be the length-normalized matrix from the Belayer alignment.

#### 2.5.3 Measuring gene trajectory similarity within and between methods

For a given gene *g*, we then calculate and plot the mean of 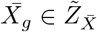 with 95% confidence interval. Additionally, we calculate Spearman’s rank correlation coefficient, *ρ*, between 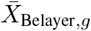 and each 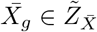 .

#### 2.5.4 Manual, ATAT, and Belayer segmentation

For all three segmentation methods of manual, ATAT, and Belayer, we utilize the manually annotated layer boundaries as a starting point. While the main purpose of our method is to generate gene expression trajectories across tissue layers, we evaluate the underlying alignment from ATAT that these trajectories are based on by quantizing the alignment into discrete segments. We use the annotated layer boundaries as an “oracle” to inform our segmentation. For each pair of consecutive layer boundaries, we take the union of path tile assignment sets for all path tiles bounded by these two layer boundaries: 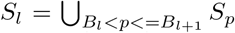 where *S*_*l*_ is the set of tiles assigned to the *l*th layer, *B*_*l*_ and *B*_*l*+1_ are the annotated boundaries on either side of layer *l, p* is a path tile, and *S*_*p*_ is the set of tiles assigned to tile *p*. Practically, this segmentation is implemented by traversing the path and marking the indices where the path crosses tiles which lie on the boundaries. For Belayer’s segmentation, we follow the tutorial presented by Ma et al. [5] and use the annotated layer boundaries and expression data as input. We then calculate the Adjusted Rand Index (ARI) to measure the similarity between manual segmentation and the segmentation derived from ATAT or Belayer.

## 3 Results

### 3.1 Biological consistency in CD and UC

We collect tiles from all 12 colon and 4 stomach samples together into a single tile dataset to train our encoder. In Figure 3, we present the path traversed between anchor points on the edge of the colonic mucosa and serosa for CD sample B and UC sample B. Our method first aligns tiles across the whole slide to the most visually similar tile along the path, thereby grouping tiles from the same tissue layer together; it accomplishes this by having learned the gradual visual transitions between tiles that are immediately adjacent to one another. As seen in Figure 3B,D, tiles from within the colonic mucosa are aligned to the tiles along the path which traverse the mucosa (spots colored in reds and yellows); other path segments similarly align their respective traversed layers to the tiles of the layers in the wider slide. However, there are some regions of spots which are not assigned according to their true layers; in the bottom-left section of Figure 3D, the region of muscularis propria is colored in the yellow-orange of the muscularis mucosa and submucosal regions. We attribute this to the visual color similarity between these regions, likely due to the faded eosin staining of this particular region of muscularis propria. However we do acknowledge this potential limitation which warrants further investigation. Ultimately we recover gene expression trajectories which correspond to existing knowledge about colonic tissue layers and these diseases.

Once we have an aligned path for a given sample, we plot known tissue and disease marker genes. Importantly, because a user has defined start and end spots in the same tissue layers, our paths and expression trajectories are anchored to the true tissue layers and can thus be meaningfully compared across multiple slides. To accomplish this, we length-normalize the expression trajectories within each sample and average the trajectories across samples of the same disease (see Section 2.4). Figure 4A-C show the averaged expression trajectories for 4 Crohn’s disease samples while Figure 4D-F show the same for 4 ulcerative colitis samples; the specific paths traversed through the CD and UC samples are shown in Figure S1 and Figure S2, respectively. The average expression trajectories for genes *EPCAM* and *ACTA2* are shown in Figure 4A,D. *EPCAM* is a known epithelial marker while *ACTA2* is expressed by smooth muscle. Because our paths across all samples traverse from the colonic mucosa, through the submucosa, and end in the muscularis propria, we would expect *EPCAM* to decrease and plateau as the path moves past the mucosa. Conversely, *ACTA2* generally increases as tissue depth increases as smooth muscle becomes more abundant towards the muscularis propria.

**Figure 4:**
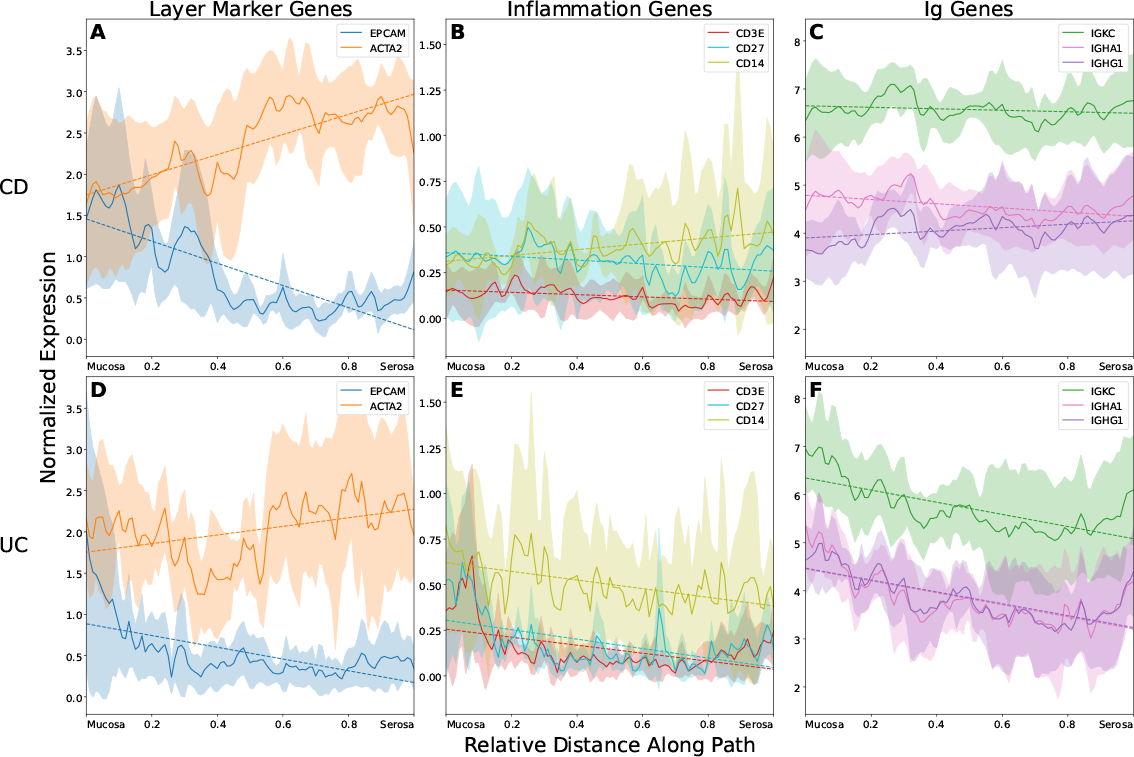
Gene expression trajectories across tissue layers for Crohn’s disease and ulcerative colitis, averaged across *N* = 4 samples per condition. For a given gene, solid lines show average trajectory with shaded region above and below indicating 1 standard deviation. Dashed lines show line of best fit for each gene’s average trajectory. For each sample, trajectories are plotted along paths spanning from the colonic mucosa to the outer serosa and are length-normalized to allow averaging between paths of differing lengths. Expression is normalized such that the average number of counts per spot across the entire slide before logarithm is 10,000. Y-axis is shared within each column. (A-C) Average trajectories from Crohn’s disease. (D-F) Average trajectories from ulcerative colitis. (A,D) Expression of known colon marker genes for intestinal epithelium (*EPCAM*) and smooth muscle (*ACTA2*). (B,E) Expression of inflammation related genes for T-cells (*CD3E*), B-cells (*CD27*), and macrophages (*CD14*). (C,F) Expression of immunoglobulin (Ig) genes for the kappa light chain (*IGKC*), IgA1 (*IGHA1*), and IgG1 (*IGHG1*).

We further plot the expression trajectories for genes associated with inflammation (Figure 4B,E) and immunoglobulins (Figure 4C,F). For the inflammation genes, we first observe that there is much greater immune cell concentration in the colonic mucosa for UC (Figure 4E) than there is for CD (Figure 4B). This is consistent with clinical and pathological knowledge that UC is restricted to the mucosal layers whereas CD is characterized by full-thickness inflammation [4]. Additionally, we observe relative enrichment of *CD14* compared to either *CD3E* or *CD27*, indicating greater macrophage activity than T-cell or B-cell activity. Again, this corroborates known disease pathology that macrophages are an important mediator of IBD [23]. Within the immunoglobulin genes, we observe similar patterns of full-thickness infiltration for CD (Figure 4C) as compared to greater mucosal involvement in UC (Figure 4F). For UC, there is a negative trend in expression of these genes as one progresses towards the serosal surface. The magnitude of expression for the selected immunoglobulin genes for the kappa-light chain, IgA1, and IgG1 is also reflective of the underlying biology. Since there are two kappa-light chains per immunoglobulin, *IGKC* has the greatest expression, followed by *IGHA1* and *IGHG1* as shown by Uzzan et al. [24]. Through this set of gene expression trajectories, we show agreement between the results of our method and known biology.

### 3.2 Internal and external method consistency

Our method relies on a user selected histological anchor points to serve as start and end point for our traversed path. We show here that the resulting gene expression trajectory is not sensitive to the choice of start and end point as we ultimately rely on the underlying histological features of the tissue. We perform this evaluation on UC sample B from Figure 3C. We first randomly sample a set of 100 reasonable start/end points on the slide, as outlined in Section 2.5.1. We then generated an alignment for each of the randomly selected start/end pairs and length-normalized them to enable comparison. See Section 2.4 for details on our length-normalization method. For each gene in the gene set selected in Figure 4, we show that the 95% confidence interval about the average trajectory is a narrow band with small variance (Figure 5A-C), demonstrating that our method is internally consistent. That is, the resulting expression trajectory within a slide is invariant to the choice of start and end point.

**Figure 5:**
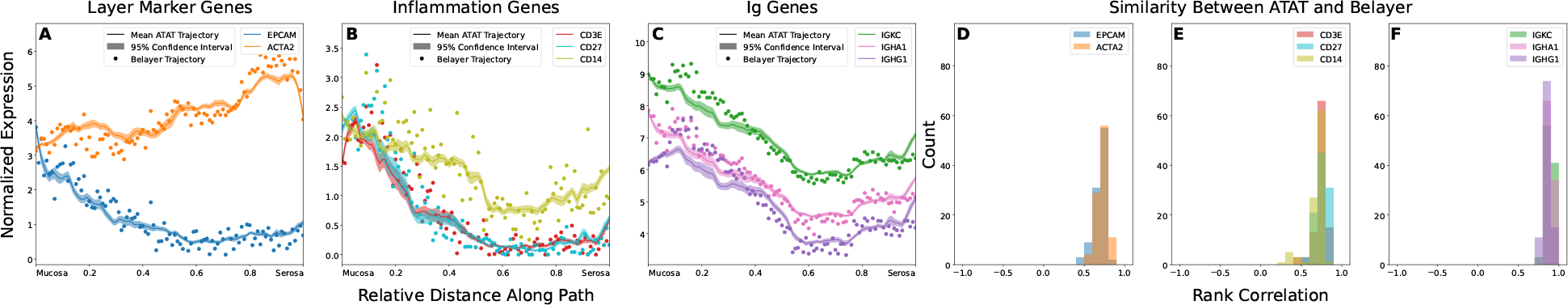
Averaged gene expression trajectories of 100 randomly sampled, yet reasonable, ATAT paths from ulcerative colitis sample B and their agreement with the trajectory derived from Belayer. (A-C) Expression trajectories derived from multiple tissue alignments of ulcerative colitis sample B. Each trajectory is length-normalized so that paths of differing lengths can be compared together. For a given gene, the solid lines show the averaged gene expression trajectories along the ATAT traversed paths of 100 randomly sampled reasonable start/end pairs with the shaded region above and below the line denoting the 95% confidence interval. The scatter plot shows the Belayer derived trajectory for the same gene. (D-F) Histogram of Spearman’s rank correlation coefficient between the Belayer derived trajectory and each ATAT derived trajectory from the randomly sampled paths. Y-axis is shared for the bottom row. (A,D), (B,E), and (C,F) show the same genes per column as in Figure 4.

In addition to demonstrating internal consistency, we show that ATAT is consistent with the state-of-the-art alignment method, Belayer [5]. We generate an alignment using Belayer for our evaluation tissue sample by manually annotating layer boundaries over the tissue sample (Section 2.5.2). Layer boundary annotations and Belayer alignments for all CD and UC samples are shown in Figure S1 and Figure S2, respectively. Similar to the randomly sampled paths, we length-normalize the Belayer alignment and plot the resulting gene expression trajectory as a scatter plot for each gene in Figure 5A-C. We quantify the similarity between ATAT and Belayer by calculating the rank correlation coefficient between the Belayer derived trajectory to each of the trajectories from the randomly sampled ATAT alignments. We plot the correlation coefficient for each gene as a histogram in Figure 5D-F. We observe a high concordance between ATAT and Belayer with almost all correlation coefficients greater than 0.5.

### 3.3 Segmenting into tissue layers

We compare the segmentation of ATAT and Belayer to manual segmentation. All three segmentation methods, of manual, ATAT, and Belayer, require as input manually annotated layer boundaries. In Figure 6, we show the segmentation results of these methods on UC sample A. We observe that while ATAT shows more noisy layers than those by manual or Belayer segmentation, ATAT is better able to capture the contours of the muscularis propria, colored in green. Across all CD and UC samples, we show that ATAT does not do significantly different from Belayer, as measured by the ARI between ATAT or Belayer and the manual segmentation (Figure 6E). We show the segmentation of all CD and UC samples in Figure S1 and Figure S2 and the table of all ARI values in Table S1.

**Figure 6:**
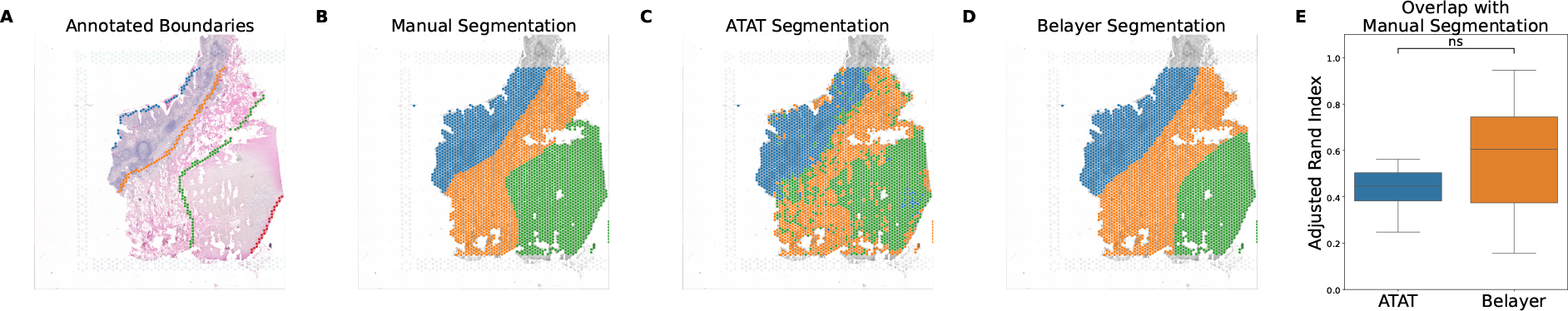
Comparison of segmentation methods on ulcerative colitis sample A, using manually annotated layer boundaries as the starting point for all segmentation methods. 3 layers are segmented using each method. Quantification of segmentation overlap is over all 8 Crohn’s disease and ulcerative colitis samples. (A) Manually annotated layer boundaries, defining 3 regions between boundaries. (B) Manual segmentation between the annotated layer boundaries. (C) ATAT Segmentation using annotated boundary crossings along the traversed path to quantize the ATAT spot assignments. (D) Belayer segmentation derived from conformal mapping of the annotated boundaries and gene expression data. (E) Adjusted Rand Index comparing the segmentation of ATAT or Belayer to the manual segmentation across all 8 samples of Crohn’s disease and ulcerative colitis. ns: not significant by independent T-test, p = 0.170.

### 3.4 Aligning external brain samples

Finally, we validate our method on external data from Maynard et al. [20] of the DLPFC. We train the encoder model on the H&E images from all 12 DLPFC samples and generate an ATAT path and alignment on sample 151673 from donor 3 (Figure 7B). On the same sample, we manually annotate layer boundaries (Figure 7A) and generate a Belayer alignment (Figure 7C). We length-normalize the resulting trajectories for ATAT and Belayer and plot in Figure 7D the gene expression for *NEFH*, a gene selected in Ma et al. [5]. We show that ATAT is able to generate a similar expression trajectory to Belayer when applied to an external dataset.

**Figure 7:**
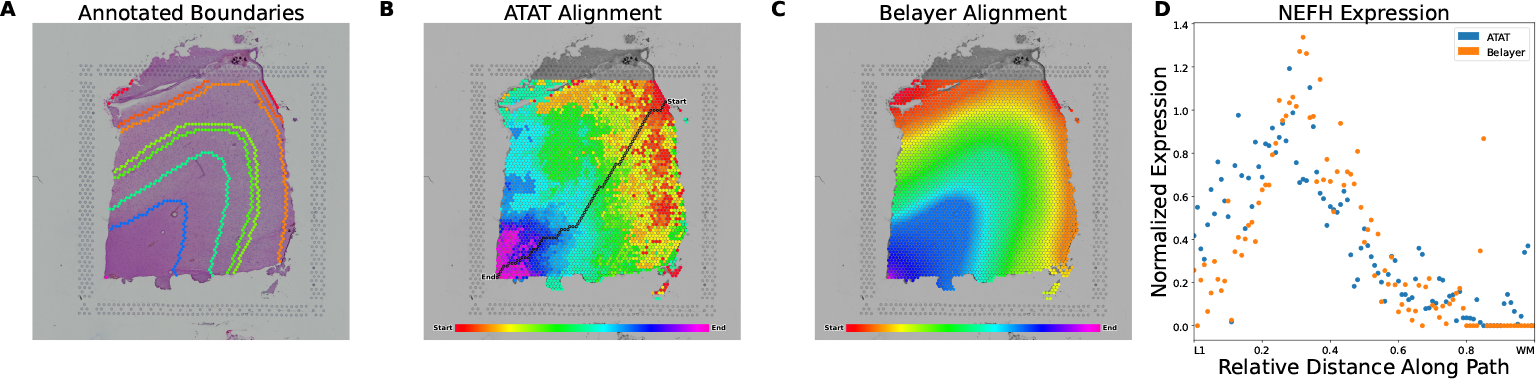
Comparison of ATAT and Belayer on alignment and expression trajectory on DLPFC tissue sample 151673. (A) Mannually annotated layer boundaries of the brain tissue layers. These annotations are based on the segmentation provided with the data. (B) ATAT traversal and alignment. (C) Belayer alignment based on manually annotated layer boundaries. (D) Length-normalized expression trajectory of *NEFH* gene generated using either ATAT alignment or Belayer alignment. Spearman’s rank correlation coefficient *ρ* = 0.725.

## 4 Discussion

In our analysis of UC and CD samples, we observed similar immune cell types driving both diseases and also identified distinguishable variations in inflammation depth between them. These results allow us to assert that ATAT is able to find biologically consistent expression patterns. Additionally, while our method requires minimal user input, we show that the resulting expression trajectory is consistent regardless of user selection when the selection successfully anchors the path to the underlying tissue histology. In our evaluation of start/end choice invariance, though we show that our method is able to generate a highly confident average trajectory when sampling many paths, user effort increases with the greater amounts of sampling required to generate many paths per sample. In our evaluation, we relied on manually annotated layer boundaries to derive the set of paths from which to sample from. We leave the decision to sample many paths per sample up to the user. This presents ATAT as a flexible, simple to use, and powerful tool that any research user can use to analyze expression patterns over histological layers.

While we emphasize our method’s ability to generate biologically relevant gene expression trajectories, many other methods aim to derive layer segmentations based on the spatial expression data [5, 6, 7, 8, 9]. Within these methods, Belayer finds the segmentation most consistent with manual segmentation [5]. As a validation of the tissue alignment generated by ATAT, tissue segmentations derived using our alignment are competitive to those found by Belayer. While segmentation using either ATAT or Belayer requires manually annotated layer boundaries, there are a plethora of deep learning based methods for tissue segmentation [25] which would likely do better than either ATAT or Belayer. An open line of investigation is whether these other models may serve as pretrained base models upon which to fine-tune for ATAT.

Nonetheless, we show that our method does not do significantly different than Belayer on segmentation. We further argue that the manual segmentation of some tissue samples is over-idealized. In our colon dataset, layer boundaries are often not sharply delineated due to disease erosions, difficulties with slide processing, or a combination of both. In the DLPFC dataset, the layers of the cortex are often difficult to distinguish and may similarly lack sharp boundaries with regions of histological overlap [26]. The segmentation based on our method may be able to more granularly define these boundaries, in both the colon and brain, as the ATAT alignment more closely models the gradual transitions between layers (Figure 7B,C, Figure S1A-F).

Additionally, in UC sample B and DLPFC sample 151673, we demonstrate that ATAT is comparable to Belayer in terms of generating biologically relevant gene expression trajectories. While Belayer is able to neatly align the ST lattice, it does so at the cost of manual annotation of all layer boundaries within the sample. If the tissue layers can be approximated by straight lines, Belayer is able to learn the alignment and layer boundaries directly from expression data without manual annotation. For UC sample B, the layer boundaries can be approximated by straight lines, however when we attempted to run Belayer with this approximation, the predicted runtime for completion on the tissue sample was over 17 days. For comparison, with manual annotation, Belayer’s alignment of spots completes in less than a minute, and its segmentation combining annotated boundaries with expression data completes on the order of tens of minutes. This tradeoff between computational time, mixing data inputs, and manual effort does not exist for our method. Our training of ATAT over 16 gastroinstestinal samples took approximately 10 hours, however multiple strategies for reducing this training time have yet to be explored, such as training for fewer epochs, reducing model parameters, or fine-tuning from pretrained models.

One additional feature of our method which we wish to highlight is the ability for ATAT to align non-linear tissue layers and complex morphologies to a path traversed through simpler, linear regions. We collected a set of examplars in Figure 8. In each of these colon CD samples, there are sections of tissue with which are not neatly stacked, highlighted by the red outline (Figure 8A-C). On these slides, the arrow annotations originating from the outlined region and ending in the non-outlined region denote matching tissue layers. For example, in Figure 8A, the epithelium and lamina propria of the colonic crypts in the center left of the slide should be aligned to the crypts in the middle of the slide; the smooth muscle in the top left should be aligned to the muscularis propria in the bottom right of the slide. ATAT is able to align these segments to a path traversed through a minimal portion of the slide, again due to the anatomical anchoring our strategy employs. With the difficult reality of preparing neatly oriented tissue samples on slides and needing to potentially fit multiple tissue samples onto a single slide, ATAT allows for the joint analysis of all tissue pieces within the slide, regardless of morphology.

**Figure 8:**
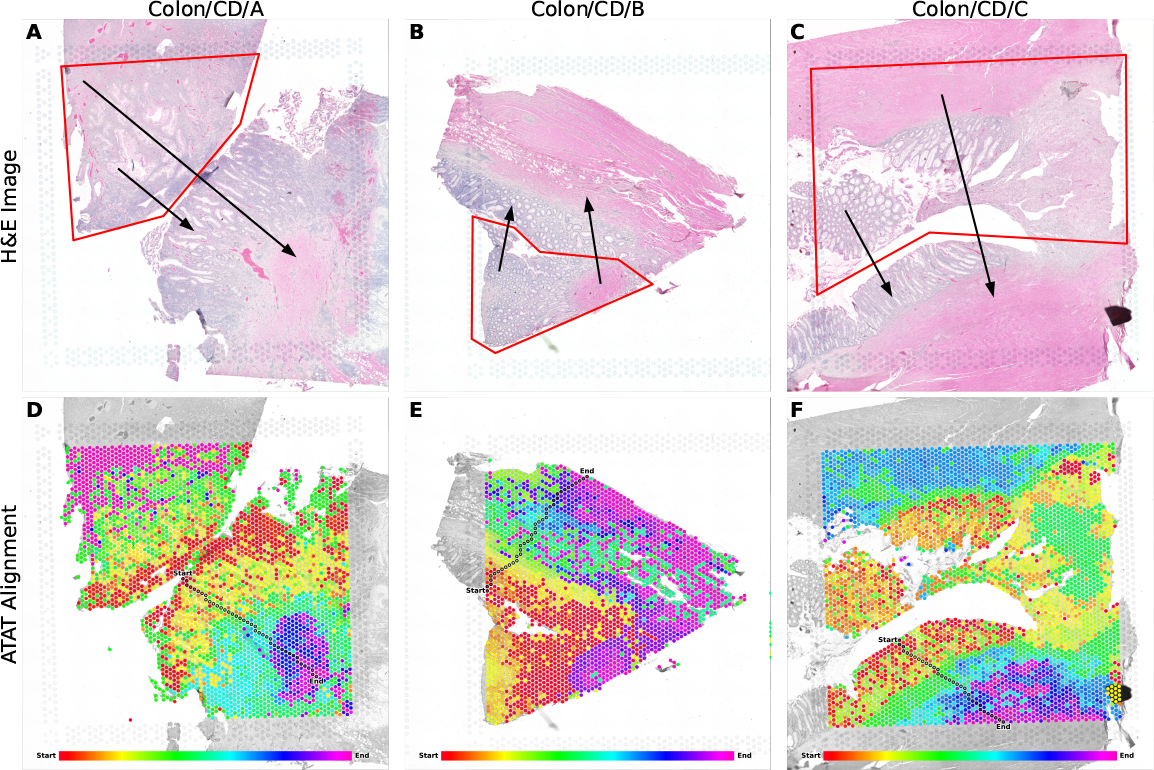
Complex tissue morphologies which are not organized in neatly stacked layers and their traversal and alignment by ATAT. (A-C) H&E images for 3 tissue samples from patients with Crohn’s disease. Regions outlined in red denote areas of tissue which are not neatly stacked, as in the non-outlined regions. Arrows between the outlined region and non-outlined region denote tissue layer similarity between the two regions. (D-F) ATAT traversal and alignment of each tissue sample.

While we present ATAT as solving multiple challenges with existing methods and working with ST data, there are potential limitations of our method as well. As discussed in Section 3.1, ATAT may result in patches of misaligned tissue. This is likely due to the visual similarity between phenotypically adjacent tissue layers and the cells which comprise them. One area in which we explored to overcome this challenge was tile augmentations, such as random flips and rotations. Yet we found the alignment to be more disorganized with such augmentations during training, likely due to the importance of the orientation of cells when learning if two tiles are adjacent or not. For example, in the inner and outer muscularis propria layers, the smooth muscle cells look otherwise similar except for their orientation depending on their layer; augmentation remove this critical piece of information for the model. While the average of expression across spots in the overall alignment is sufficient, based on internal and external consistency evaluations, achieving more perfect alignment is an open task. Finally, while we learn spatially aware tile representations in a self-supervised manner, we ultimately do still require minimal user input to define the start and end spots upon which to anchor our path. We argue that this is a reasonable tradeoff between manual effort and user expertise.

We propose ATAT as a novel tool to enable biological discovery by facilitating the exploration of ST data. The traversal and alignment of samples and the derivation of expression trajectories by ATAT offers a biologically relevant dimensionality reduction for data exploration. Our analyses confirmed that our method consistently yields anticipated outcomes for diseases and tissues with recognized spatial phenotypes. As the number of spatial transcriptomic sequencing libraries grows, tools such as ATAT are necessary to automate much of the data exploration with minimal user intervention. Thus, ATAT presents a powerful histological imaging based machine learning method which enables access to crucial molecular and cellular data in tissue biopsies and minimizes manual annotation in spatial transcriptomics analysis.

## 5 Acknowledgements

We thank the participants of our study whose data enabled us to develop ATAT. The authors thank Elizabeth Moison, Hugh Yeh, and Salvador Norton de Matos for insightful discussions. This work was partially supported by NIH DP2 NIAID New Innovator Award (DP2AI177884), the University of Chicago’s Center for Interdisciplinary Study of Inflammatory Intestinal Disorders (C-IID) (P30 DK42086), the Institute for Translational Medicine (ITM) (5UL1TR002389-05), and a Hardware Grant from NVIDIA.

## 6 Code and data availability

The code for ATAT can be found at https://github.com/StevenSong/tissue-alignment. The repository includes code and documentation for training the vision encoder model and for path traversal and alignment. Data for colon and stomach tissue samples are available upon reasonable request to the corresponding author.

