## Supplemental Materials for "ATAT: Automated Tissue Alignment and Traversal in Spatial Transcriptomics with Self-Supervised Learning"

#### A Crohn's Disease Alignment

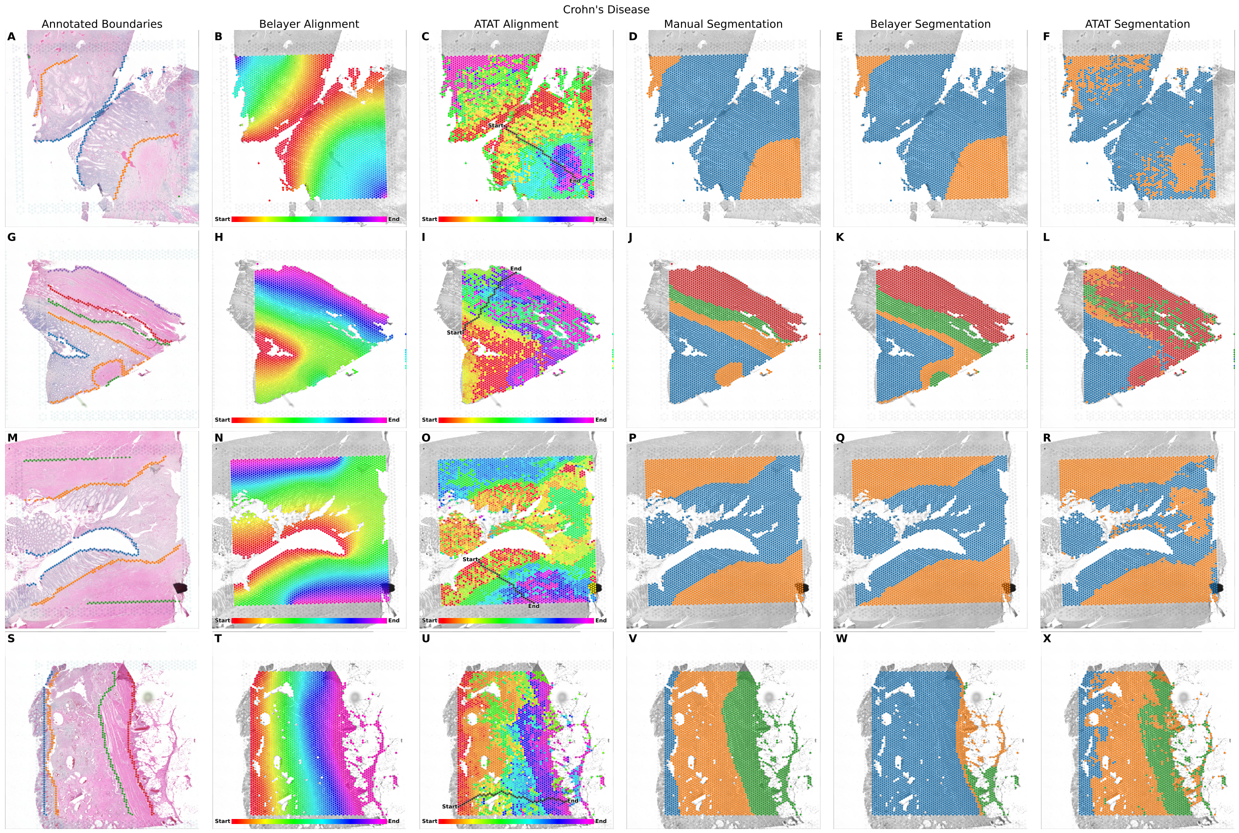


Figure S1: All annotations, alignments, and segmentations of Crohn's disease samples. (A-F) Crohn's disease sample A. (G-L) Crohn's disease sample B. (M-R) Crohn's disease sample C. (S-X) Crohn's disease sample D. (A,G,M,S) Manually annotated layer boundaries. Layer boundaries may be limited by the extent of the hexagonal grid within the capture area of the ST slide. (B,H,N,T) Belayer alignment. The hexagonal grid of spots is colored according to the depth scalar assigned to each spot by Belayer. (C,I,O,U) ATAT alignment between user selected anchor points. Spots colored according to the most visually similar spot along the traversed path. (D,J,P,V) Manual segmentation based on annotated boundaries. (E,K,Q,W) Belayer segmentation based on Belayer alignment and expression data. (F,L,R,X) ATAT segmentation. Spot assignments are quantized using crossings over the annotated boundaries along the path.

### B Ulcerative Colitis Alignment

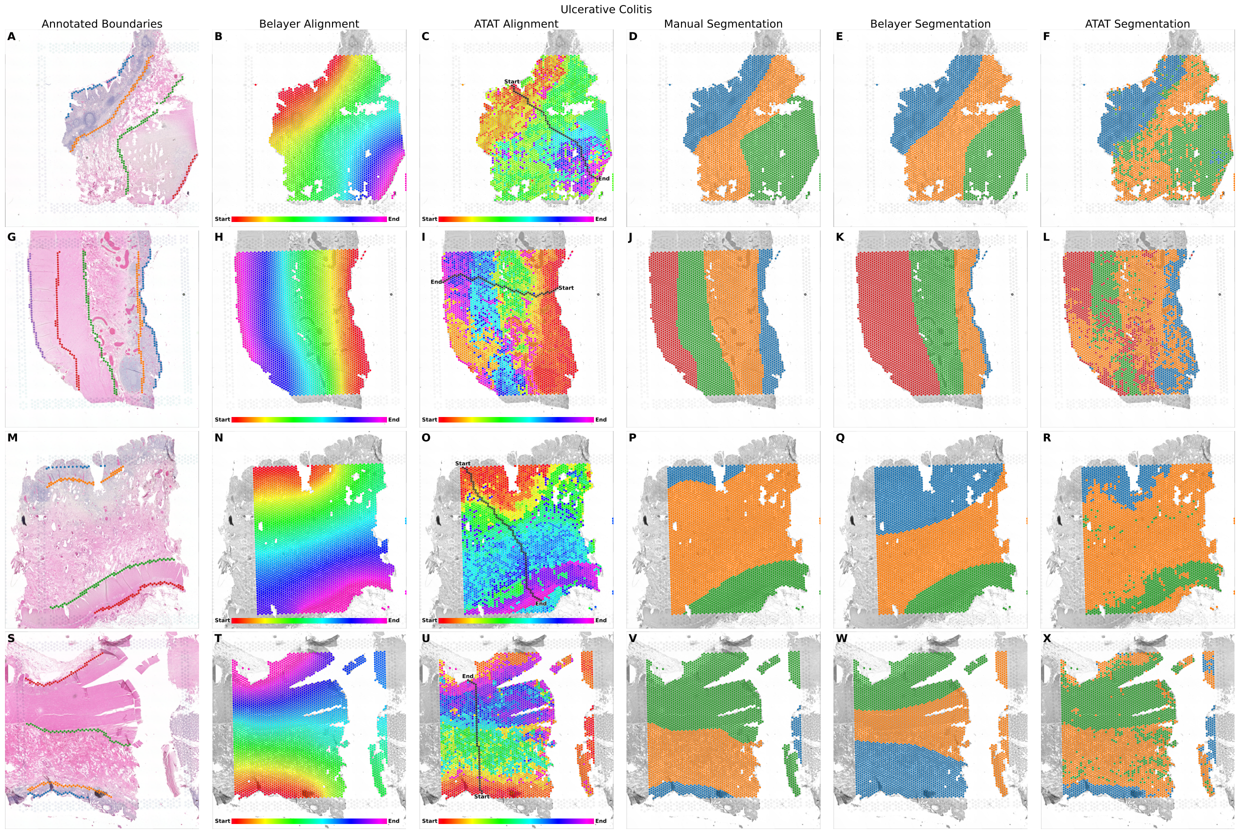


Figure S2: All annotations, alignments, and segmentations of ulcerative colitis samples. Columns are the same as in Figure S1 but for UC samples; refer to the caption of the CD Figure for more details on each column. (A-F) Ulcerative colitis sample A. (G-L) Ulcerative colitis sample B. (M-R) Ulcerative colitis sample C. (S-X) Ulcerative colitis sample D.

#### C Reasonable Path Sampling

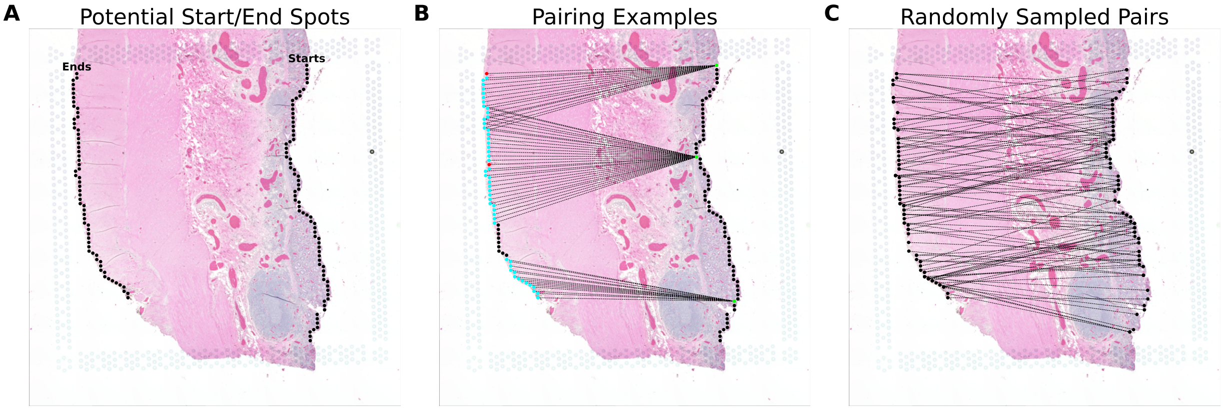


Figure S3: Path sampling method for evaluation of ATAT using UC sample B. (A) The set of potential start and end spots along the right and left edge, respectively, of the tissue sample. (B) Selected examples of reasonable start and end pairings that a user might make. Initially, each green start spot is paired with a red end spot by matching spots by index in either list of spots sorted by y-coordinate, relative to the top-most spot. For a given green start spot, its set of paired end spots is then increased by taking teal end spots whose index in the end list lie within a radius of 15 spots on either side of the paired red end spot. For a given green start spot which does not have a corresponding red end spot, we utilize a virtual red end spot from which to consider expanded teal end spots, as in the bottom-most exemplar green spot. For UC sample B, there are $1,796$ reasonable start/end pairs. (C) 100 start/end pairs randomly sampled from the set of all reasonable start/end pairs.

### D Quantification of Segmentation Comparison

| Sample | | ARI with Manual | |
|---|---|---|---|
| Disease | Section | ATAT | Belayer |
| CD | A | 0.310 | 0.946 |
| CD | B | 0.456 | 0.721 |
| CD | C | 0.562 | 0.813 |
| CD | D | 0.500 | 0.156 |
| UC | A | 0.437 | 0.655 |
| UC | B | 0.248 | 0.555 |
| UC | C | 0.515 | 0.376 |
| UC | D | 0.406 | 0.366 |

Table S1: Adjusted Rand Index for each colon IBD sample comparing either ATAT or Belayer segmentation with manual segmentation.
